# Extensive Phenomenological Overlap between Induced and Naturally-Occurring Synaesthetic Experiences

**DOI:** 10.1101/2020.08.03.228692

**Authors:** David. J. Schwartzman, Ales Oblak, Nicolas Rothen, Daniel Bor, Anil. K. Seth

## Abstract

Grapheme-colour synaesthesia (GCS) is defined by additional perceptual experiences, which are automatically and consistently triggered by specific inducing stimuli. The associative nature of GCS has motivated attempts to induce synaesthesia by means of associative learning. Two recent studies have shown that extensive associative training can generate not only behavioural (consistency and automaticity) and neurophysiological markers of GCS, but also synaesthesia-like phenomenology [1,2]. However, these studies provided only superficial descriptions regarding the training-related changes in subjective experience: they did not directly assess how closely induced synaesthetic experiences mirror those found in natural GCS. Here we report an extended qualitative analysis of the transcripts of the semi-structured interviews obtained following the completion of the associative training protocol used by [2]. In addition, we performed a comparable analysis of responses to an interview with a new population of natural occurring grapheme-colour synaesthetes (NOS), allowing us to directly compare the phenomenological dimensions of induced and naturally occurring synaesthetic experience. Our results provide an extensive addition to the description of the phenomenology of NOS experience, revealing a high degree of heterogeneity both within and across all experiential categories. Capitalising on this unique level of detail, we identified a number of shared experiential categories between NOS and induced synaesthesia-like (ISL) groups, including: *stability of experience, location of colour experience, shape of co-occurring colour experience, relative strength of colour experience and automaticity of colour experience*. Only the automaticity of colour experience differed significantly between the two groups: NOS experience was reported as being mostly automatic, whereas induced ISL were mostly described as being ‘wilful’. We observed three additional experiential categories relating to the automaticity of synaesthetic experience within the NOS group: *contextually varied experience, semi-automatic experience* and *reflective association*, which suggests that, as with other experiential categories, the automaticity of synaesthetic experience is also highly heterogeneous. Our results provide new evidence that that intensive training of letter-colour associations can alter conscious perceptual experiences in non-synaesthetes, and that such alterations produce synaesthesia-like phenomenology which substantially resembles similarities to natural grapheme-colour synaesthesia.

## 1.0 Introduction

In grapheme-colour synaesthesia (GCS), the presence of achromatic (black) letters (inducer) triggers the experience of colour (concurrent). The defining characteristics of GCS include the automaticity of the concurrent experience (i.e., not experienced as being under voluntary control), normally assessed using an adapted version of the Stroop task [3,4,5], and the consistency of the associations (i.e., repeated presentations of an inducer will elicit highly similar concurrent experiences)[6,7,8]. Typically, natural synaesthetes differ in the nature of their synesthetic experiences. Some grapheme-colour synaesthetes, referred to as *projectors*, experience their concurrent outside bodily space, ‘projected’ into the world. Others, referred to as *associators*, report that their experiences exist within their internal mental space without any distinct spatiality [9,10,11]. A less frequently noted characteristic of GCS is that concurrent experiences usually lack what is called *perceptual presence* [12]. This means that a concurrent experience is not usually confused with, or perceived as, a ‘really-existing’ property of the world. In GCS, even though an inducer elicits an additional colour experience (e.g., red), grapheme-colour synaesthetes will still perceive the inducer as being the colour it actually is (e.g., black).

The associative nature of GCS has led researchers to theorise that a learning component must be involved in the development of synaesthetic associations (for review see [13]). This has prompted a number of studies to investigate if it is possible to train non-synaesthetic (neurotypical) individuals to have synaesthesia-like experiences [1,2,4,14,15,16]. In two previous studies of this kind, we used extensive and adaptive associative training regimes towards this goal [1,2]. Using the gold-standard consistency test (Eagleman, Kagan, Nelson, Sagaram, & Sarma, 2007; Rothen et al., 2013; www.synesthete.org), we found in both studies, that performance for trained letter-colour pairs passed the threshold indicative of synaesthetic experience. In addition, participants also displayed synaesthesia-like behaviour for trained letters on a synaesthetic equivalent of the Stroop test (demonstrated by greater interference effects and slower response times in incongruent trials) [4,5]. Critically, the majority of participants in the first study [1], and all participants in the second study [2] self-reported phenomenology suggestive of natural synaesthesia. However, these studies only provided relatively superficial descriptions of these training-induced changes in phenomenology.

In this study, we investigate in more detail the degree to which trained and natural synaesthetic experiences are similar. We report an in-depth qualitative analysis of the transcripts of the semi-structured interview obtained following the completion of the training protocol (data taken from [2]). We accompany this analysis with an equivalent analysis performed on interview transcripts obtained from similar semi-structured interviews obtained from a new sample of natural GCS subjects. Comparing these detailed phenomenological datasets, we provide a direct comparison between these two types of unusual perceptual experience.

## 2.0 Material and Methods

### 2.1 Participants

The Induced Synaesthesia-Like (ISL) group consisted of 18 non-synesthetes (15 women, mean age = 23, SD = 3.08), whose data and interview recordings were taken from [2]. The Naturally Occurring Synaesthesia (NOS) group consisted of 15 grapheme-colour synaesthetes (14 women, mean age = 43.3, SD = 11.43). Experiments were undertaken with the understanding and written consent of each ISL participant. All NOS participants were recruited from the University of Sussex synaesthesia database based on indicative consistency scores of GCS (www.synesthete.org; [7]). Informed consent was obtained from NOS participants prior to the beginning of the interview. The experiment was approved by the University of Sussex ethics committee.

### 2.2 Behavioural tests

All participants completed the internet-based standardized grapheme-colour consistency test (www.synesthete.org; [7]). The test presents each participant with the graphemes A–Z three times in randomized order, for each presentation of a letter participants were asked to select the colour that best fit with each grapheme. The ISL group performed this test twice, once before and once upon completion of the training paradigm [2]. As part of their inclusion in the University of Sussex’s synaesthesia database, all NOS participants completed the grapheme-colour consistency test (once).

### 2.3 Phenomenological analysis

Our research design was constructed around three related approaches in empirical phenomenology and qualitative research: two-phase research [17], sequential analysis [18] and theoretical sampling [19]. Based on these approaches, in Phase one, we gathered data from the ISL group that aimed at constraining our object of inquiry, so that in Phase two, we could acquire more precise and focused qualitative data from NOS participants. The order of the Phases was based on the historical dependencies between the two studies. The semi-structured interview used in [2] was carried out first and enquired about the core characteristics of synaesthesia (e.g., consistency, automaticity, unidirectionality etc.), which based on empirical evidence from NOS studies. This meant that the structure of the interview used in Phase two (with NOS participants) was very similar to that used in Rothen et al., (2018). Specifically, in the first Phase of the study, we conducted a semi-structured interview on participants from our training study [2]. Semi-structured interviews are particularly well-suited for investigating difficult-to-define experiences[2]. The manner in which the participant speaks and phrases particular responses can lead to a greater understanding of participant’s experiences, while still allowing the participant the freedom to elaborate [20]. Based on ISL participant’s subjective reports, we designed the second Phase of research, in which we gathered commensurable reports from individuals with naturally occurring GCS (the NOS group). In the following subsections, we present the protocol for each Phase of research.

### 2.4 Phase 1: Subjective reports of induced synaesthesia-like experiences

In the first Phase, following the completion of a 5-week training battery, all (ISL) participants performed a semi-structured interview designed to assess their perceptual phenomenology during exposure to 13 achromatic trained letters (for details see [2]). Due to the nature of our research questions, only sections of the interview that explicitly asked participants to describe their colour experiences were transcribed. These sections included responses to the question “Look at this page that has the 13 letters you have been trained over the last 5 weeks, to associate with 13 specific colours. I want you to describe any associated colour experience you have when looking at these letters”. Additionally, within the interview a subset of 14 participants were asked to compare the strength or vividness of their strongest trained synaesthesia-like colour experience (e.g. ‘r’ is red) to a life-long colour association of a real-world object (e.g. the specific shade of red associated with an English post-box).

Further, we constrained the focus of responses in a funnel-like structure frequently used in semi-structured interviews[21], by including responses and discussion to a forced choice question that required participants to localise their trained synaesthesia-like experiences in space. Question: “Which statement characterises your grapheme-colour associations best?

Whenever I see a letter…

- There is only that letter, but no colour at all. I can’t even think of an associated colour, no matter how hard I try.

-I know the associated colour, but I never see it.

-I see the colour in front of my mind’s eye.

-I see the colour outside my head (i.e., a few inches away).

-I see the colour floating on the surface wherever the letter is.”

The interviews were transcribed following established methods of analysing subjective reports [22,23]. The analysis of the transcripts was carried out in two stages. First, the description of actual subjective experiences relevant to our line of enquiry were highlighted within the transcripts, while other descriptions (such as clarification about the question, or irrelevant discussion) were removed from the transcripts. We then performed a content analysis on the transcripts for all participants (N = 18) (transcripts are available https://osf.io/e367d/?view_only=dd61d42daa7a4c848023b89bd38789f8). Experiences within the transcripts were then classified with respect to their specific content. We avoided using preconceived categories, instead allowing the categories and names for categories to emerge from the data [24].

The data was then analysed according to the principles of *content analysis*, whereby codes are not assigned to the data based on the concrete words used, but rather according to the underlying meaning [25]. We used inductive coding, meaning that we ascribed abstract, more general description to the raw data without recourse to established theoretical constructs [cf. 19]. This approach was chosen as it recognises that the knowledge gathered through interviews is jointly constructed by the researcher and participant, rather than being discovered in an observer-independent fashion, while still providing an established method for the analysis of qualitative data [26,27]. Following the logic of induction (i.e., moving from concrete raw data to more abstract, general descriptions), we assigned descriptive tags to the transcribed interviews. In doing so, we did not follow a pre-existing framework. Rather, we allowed experiential categories to emerge naturally from the experiential reports. Afterwards, we grouped specific codes together based on their structural (i.e., descriptive) similarities. Note that our coding system, while open-ended, was not assumption-free. We constructed a taxonomy of experiential categories such that it allowed for comparison of phenomenological dimensions of induced synaesthesia-like with naturally-occurring synaesthetic experiences. The validity of data acquisition and analysis was evaluated using two methods: a) *the annotated codebook* and b) *intercoder verification*. In qualitative research, the annotated codebook approach [28] is an instrument that serves three purposes. Firstly, it is a way of organizing qualitative data by establishing meaningful relationships between individual reports. Secondly, a fully specified codebook (i.e., a codebook in which meaningful relationships between all the categories is established) provides a method for establishing that the gathered data is ‘deep’ enough for theory-construction [29,30]. Thirdly, it represents an instrument in which the coding process is described in sufficient detail to enable independent replication. The annotated codebook itself consists of two instruments: the saturation grid and the codebook itself. The saturation grid is a tabulation in which for each participant we note new instances of codes established from the interviews. Once we observe no new categories for an individual, conceptual depth has been achieved (also called saturation), and it is no longer necessary to conduct further interviews [31]. Table 1 displays a condensed version of the saturation grid for the ISL group (a fully specified saturation grid for the ISL group can be found in the supplementary materials). As can be seen in Table 1, conceptual depth was reached with participant 17: after this participant, no new codes were identified.

**Table 1.** Condensed version of a sample saturation grid for the ISL group. Only participants included in [2] were used for analysis.

|  |  |  |  |  |  |  |  |  |  |  |  |  |  |  |  |  |  |  |
| --- | --- | --- | --- | --- | --- | --- | --- | --- | --- | --- | --- | --- | --- | --- | --- | --- | --- | --- |
| Participants | 1 | 2 | 3 | 4 | 5 | 6 | 7 | 9 | 11 | 12 | 13 | 14 | 16 | 17 | 18 | 20 | 21 | 22 |
| New codes | 9 | 3 | 1 | 2 | 0 | 1 | 1 | 0 | 0 | 0 | 1 | 0 | 0 | 1 | 0 | 0 | 0 | 0 |

The codebook was constructed according to standard approaches in empirical phenomenology [e.g., 32–34]. Each entry in the codebook is specified according to the following elements: name, description, subcategories, examples, and considerations. Considerations describe any specific differences between similar categories as well as concerns that may have emerged during the coding process (see supplementary materials for the full annotated codebooks https://osf.io/e367d/?view_only=dd61d42daa7a4c848023b89bd38789f8). A second measure that was used to ensure the validity of the analysis was intercoder verification. Two coders (DS and AO) coded all transcripts independently and then compared their results for consistency [35]. Experiential categories were only identified and carried forward for analysis if identified by both coders.

### 2.5 Phase 2: Subjective reports of naturally-occurring synaesthetic experiences

In the second Phase of the study we collected additional data using a semi-structured interview from individuals with NOS. The interviewing protocol for the second Phase was constructed based on the findings from the first Phase; i.e., the interview guided participants towards observing and reporting on the main phenomenological dimensions that we observed in the transcripts from ISL experiences. Specifically, whenever the participants used language based on folk psychological theories or specific theories of synaesthesia, the interviewer used precise follow-up questions prompting the participants to describe their experience in more concrete terms. The interviews were conducted via video conference (Skype or WhatsApp). Audio was recorded using a dictaphone; video was not recorded.

The semi-structured interview followed a tripartite structure. First, we obtained general information about the nature of the participants’ synaesthetic experiences, including questions relevant to the situational demands of the interview:

⍰ Have your synaesthetic experiences always been stable? Have they changed throughout the course of your life?
⍰ Do your synaesthetic experiences change depending on your mood, time of day, your level of fatigue?
⍰ Do your synaesthetic experiences change depending on whether you are reading text from a computer screen or a piece of paper?

We also collected demographic information and whether participants were professionally involved with academic mind-science disciplines, which may have led to theoretically-laden description of experience (e.g. Psychology, Neuroscience). The second Phase also involved participants viewing 13 graphemes, which were presented (via screen-share) individually in the same size and font as used in our training study [2], and they were asked to report on their associated colour experience. Following the presentation of each grapheme, they were asked to verbally identify the RGB code that best reflected their specific colour experience in relation to the grapheme, using an online colour picker (https://htmlcolorcodes.com/color-picker/). In this Phase, the participants also referred to the strength of the association between the grapheme and the colour on a scale from 1 to 10, with the weakest association being 1 and the strongest being 10. This section of the interview concluded with the participants having to report on the overall automaticity of their synaesthetic experience on a scale from 1 to 10, with 10 being fully automatic.

The third section of the interview examined participant’s synaesthetic phenomenology in more detail by using a number of open-ended questions, as well as a specific mental exercise.

The open-ended questions were:

⍰ Can you describe the details of the colour experience (shape, letter, block etc.; precise location your synaesthetic experiences take place).
⍰ When looking at a letter, for instance, ‘r’, are you aware of both the actual colour of the letter on the page (black) and your synaesthetic colour experience? Or does one overlay the other?
⍰ Can you describe how your synaesthetic colour experience are similar/not similar to the colour of a real-world object?
⍰ Do your synaesthetic experiences change if the grapheme is capitalized or in a different font?
⍰ Are there any other types of experiences you have had in your life, which are similar to your synaesthetic experiences?

The mental exercise was:

⍰ Can you bring to mind a typical everyday colour association, such as *post-boxes are red* or *the sky is blue*. How is the experience of this colour association similar or different to your synaesthetic associations?

Finally, we asked the same forced choice question as with the ISL group, which required participants to localise their colour experience in space i.e. “Which statement characterises your grapheme-colour associations best?” Forced choice question options were the same as for the ISL group.

The audio recordings of the interviews were transcribed verbatim. Whenever participants referred to colour shades that were unfamiliar to the researcher by name, a picture of that colour was included in the transcript. If the participants referred to phonemes rather than graphemes, the audio in question was transcribed using the International Phonetic Alphabet. Subsequently, the transcripts from the NOS group were analysed using the same method as for the ISL group [18].

The validity of the data in the second Phase was established using the same methods as for Phase 1, through the construction of an annotated codebook and intercoder verification (DS and AO). Finally, as with ISL experience, we determined conceptual depth by constructing a saturation grid (Table 2), which revealed that we had reached saturation by participant NOS-10.

**Table 2.** Condensed version of saturation grid for the NOS group.

| NOS Participant | 1 | 2 | 3 | 4 | 5 | 6 | 7 | 8 | 9 | 10 | 11 | 12 | 13 | 14 | 15 |
| --- | --- | --- | --- | --- | --- | --- | --- | --- | --- | --- | --- | --- | --- | --- | --- |
| New codes | 11 | 10 | 2 | 1 | 1 | 3 | 2 | 0 | 1 | 1 | 0 | 0 | 0 | 0 | 0 |

Finally, a chi-square test of independence was performed to examine the relation between ISL and NOS groups for each experiential category. We only included commensurate values and excluded instances where no values could be induced from the raw data. This statistical comparison for each experiential category should be viewed as additional information to aid in the interpretation of qualitative findings, rather than a rigorous quantitative analysis.

## 3.0 Results and Discussion

In this study we applied a qualitative analysis of interview transcripts to provide a detailed description of both training Induced Synaesthesia-Like **(ISL)** and Naturally Occurring Synaesthesia **(NOS)** experiences, enabling us to directly compare these two types of perceptual experience. Our phenomenological analysis of the transcripts from both groups identified six main overlapping categories of experience. Transcripts of interviews as well as the annotated codebooks are available online https://osf.io/e367d/?view_only=dd61d42daa7a4c848023b89bd38789f8.

Responses to questions about the nature of the NOS participants’ synaesthetic experiences revealed that six out of the fifteen participants worked in mind-science areas, raising the possibility that their reports were informed by their theoretical knowledge about synaesthesia. Verifying that the situational context of the interview (video conference) did not affect their concurrent experience, responses to the question ‘Do your synaesthetic experiences change depending on whether you are reading a text from a computer screen or a piece of paper?’, revealed that the synaesthetic experiences of all participants (N=15) were not affected by the medium on which the grapheme was presented.

As described in our previous study [2], all ISL participants reported synesthetic phenomenology following completion of the training regime. Results of the grapheme-colour consistency test (www.synesthete.org; [7]) revealed that post-training consistency scores were on average below the established threshold value of 135 in CIELUV colour space, a level indicative of natural GCS (see top panel of Fig. 3C in [2]). Consistency results for the same test verified that all NOS participants were also below the threshold indicative of natural GCS (M = 76.53; SD = 21). In addition, all ISL participants also demonstrated letter-specific behavioural effects indicative of automaticity (e.g., synaesthetic Stroop interference effect), which was further supported by reports that their synaesthesia-like experiences were automatic (assessed via forced-choice question).

### 3.1 Identified phenomenological dimensions

The main phenomenological dimensions that were constructed from the content analysis of the raw transcripts (verified by both coders) for the ISL group, which were used to compare and later observed in the NOS group were:

⍰ *Stability of colour experience*. This category describes the degree of within-subject variation in colour experience for each letter, including its strength and automaticity.
⍰ *Location of colour experience*. Within the transcripts we identified two levels of description referring to the location of the colour experience: the first, *location*, describes whether the colour experience takes place within the participants’ mental space or is externally localised. The second level of description is *location* (*specified*), describes where in relation to the inducing stimulus the colour experience occurs.
⍰ *Shape of colour experience*. This category describes differences in the visual quality of the induced synaesthesia-like experience. In natural GCS variations exist in the precise visual qualities of the colour experience [36,37], such as discernible geometric shapes, letters, auras or as a totality of colour.
⍰ *Relative strength of colour experience*. This category refers to a specific mental exercise participants were asked to perform during the interview. Participants were asked to bring to mind a strong real-world colour association (for example, the specific shade of red of an English post box). Importantly, the participants had to be aware of this association as something that remained stable throughout their lives. Then, during the mental exercise they were asked to compare their associated colour experience with this real-world colour association, and report how the two experiences compared to each other in terms of the strength and vividness of experience.
⍰ *Automaticity of colour experience*. This category refers to how automatically participants colour experience occurred. On the broadest level, automaticity refers to the distinction between colour associations being experienced as something that the participants need to actively invoke (coded as *willful*) and something that happens to them without any mental effort (coded as *automatic*).

### 3.2 Comparisons of experiential categories between ISL and NOS groups

We now discuss the results of each experiential category, their relative similarities and differences, and the continuity between these reports and those found in the synaesthesia literature:

#### 3.2.1 Stability of colour experience

This category describes a high-order description of induced-synaesthetic experience in terms of how varied the experience was for each participant across all presented graphemes. It encompasses aspects of experience, such as the vividness of the associated colour experience and how effortful it was to bring the colour experience into awareness and how these aspects of experience varied from grapheme to grapheme. We highlight that the experiential categories, *location, shape* and *automaticity of colour experience* may also be evaluated with regards to *stability of colour experience*, but within the context of the interviews these categories were identified as separate dimensions of experience.

As is common in natural synaesthesia[37,38], the majority of ISL participants reported that their experience was *heterogeneous* (15 out of 17), displaying a high degree of variability in the strength and vividness of their synaesthesia-like colour experience associated with each letter i.e. in general, the letter ‘r’ produced a more vivid colour experience (red) than the letter ‘u’ (grey). Only two ISL participants reported a *homogenous* colour experience for all graphemes. For some graphemes, the synaesthesia-like colour association was reported as being automatic, whereas for other graphemes, it required mental effort to experience the associated colour (see Table 4).

We found a similar degree of variability within the phenomenological space of NOS participants. Ten NOS participants reported (N = 15) that their concurrent experiences were highly varied in terms of strength and automaticity between individual graphemes (coded as *heterogeneous*), with only 5 participants reporting that their experiences were homogenous across all graphemes (see Table 4). A chi-square test of independence, found no significant difference between the ISL and NOS groups for this experiential category: χ2(1, N = 32) = 1.091, *p* = 0.296. Cramér’s V measure of association, ϕc, was 0.185. In terms of the stability of synaesthetic experiences throughout their lives, eight NOS participants reported that their synaesthetic experiences had remained stable throughout their life. Four participants reported that they observed minor changes in synaesthetic experiences throughout their lives, including changes in the synaesthetic colour associated with single letters or words (e.g., NOS-02 reports the word for *Saturday* shifted from blue to grey at the onset of adulthood), or variability in the intensity of synaesthetic colours (e.g., both NOS-11 and NOS-04 report that they sometimes worry that they may have “lost their colours”). Three participants reported more major changes in their synaesthetic experiences, describing that all of their synaesthetic colours had become faded with age [cf. also 39]. The second dimension of stability of synaesthetic experience is whether the colour of their synaesthetic experience changes with context. Eight participants (N = 15) reported that their colours are present even during goal-oriented behaviour, such as reading. Conversely, seven participants reported that when reading, they were able to push their synaesthetic experiences out of their awareness.

Previous research has shown that the letter–colour associations of synaesthetes are not random, but in fact display some structure [40]. In both of our previous training studies, the letter-colour associations chosen for training were based on their prevalence in synaesthetic and non-synaesthetic populations. Within natural synaesthesia, the majority of these letter–colour associations displayed a clear semantic association (e.g. r = red)(see Fig 3, [40]; also [1]). In our first training study, we found that even before training, letter-colour pairings that exhibited a strong semantic association caused a significant Stroop interference effect. Following training, Stroop interference effects increased, but were almost entirely driven by those letters that exhibited a clear semantic association [Fig 1, 1]. In addition, we found that synaesthesia-like colour experiences were particularly strong for those letter-colour associations that involved the same semantic component [1]. These observations raise the possibility that conceptual associations may facilitate the development of at least some letter–colour associations in natural synaesthesia, which in turn, may explain why they were so successful in driving synaesthesia-like behaviours and phenomenology in our training studies.

#### 3.2.2 Location of colour experience

Within this category we constructed two levels of description. The higher-order level of description (referred to as *location*) describes whether the participants colour experience was located within their mental space or if it was externally localised. The lower-order level of description (*location* (*specified*)) describes the location of the colour experience in relation to the grapheme: *letter-adjacent*, denotes situations in which the colour was experienced proximal to the letter; conversely, *letter-overlapping* describes the associated colour experience occurring precisely over the letter.

##### Location

For the category *location* we found that, as reported in NOS [41,42], the ISL group differed substantially in the description of the location of their synaesthesia-like experience. Some (9 out of 17 ISL participants) reported their synaesthesia-like experience as occurring in external space (*projector-like*), whereas others (8 out of 17 ISL participants) reported them as occurring within their “mind’s eye” (*associator-like*). Responses to the forced choice question: “Which statement characterises your grapheme-colour associations best?” revealed a slightly different picture (see Table 3), which may be due to the constrained nature of the forced choice questioning. 12 out of 18 ISL participants reporting their synaesthesia-like experience occurred in front of their “mind’s eye”, and the rest reporting that they occurred ‘floating on the surface wherever the letter is’.

**Table 3.** Absolute frequencies of forced-choice responses to the question “Which statement characterises your grapheme-colour associations best?” for ISL and NOS groups.

| Forced choice statement | ISL:<br>Responses | NOS: Primary<br>response | NOS: Secondary<br>response |
| --- | --- | --- | --- |
| Whenever I see a letter, there is only that letter, but no colour at all. I can't even think of an associated colour, no matter how hard I try. | 0 | 0 | 0 |
| Whenever I see a letter, I know the associated colour, but I never see it. | 0 | 2 | 2 |
| Whenever I see a letter, I see the colour in front of my mind's eye. | 12 | 11 | 0 |
| Whenever I see a letter, I see the colour outside my head (i.e., a few inches away). | 0 | 0 | 2 |
| Whenever I see a letter, I see the colour floating on the surface wherever the letter is. | 6 | 2 | 4 |

Within the NOS group, we found fewer projector-like descriptions (3 out of 15), with the majority (12 out of 15) describing their concurrent experience as occurring within their *mental space* (see Table 4). This finding was supported by responses to a forced choice question that required participants to localise their synaesthetic experiences in space (see Table 3). As responses to the forced choice question were taken in the context of a face-to-face interview, NOS participants had the opportunity to elaborate on their answers. The table therefore contains two counts: *primary* and *secondary features* for individuals with NOS. Primary feature represents the count of their typical experiences, whereas the secondary feature represents experiences that may happen (usually qualified in the interviews with phrases “It happens sometimes,” or “It happens in specific circumstances”).

**Table 4.** Frequency distribution of phenomenological dimensions associated with ISL and NOS experience. The symbol “/” represents values that were not induced from the data in that specific group.

| Phenomenological dimensions of ISL and NOS experience |  |  |  |  |  |  |  |
| --- | --- | --- | --- | --- | --- | --- | --- |
| Stability of experience | Value | Heterogeneous | Homogenous | NA |  |  |  |
|  | f(ISL) | 15 | 2 | 1 |  |  |  |
|  | f(NOS) | 10 | 5 | 0 |  |  |  |
| Strength of association | Value | Weaker than colour association | Equal to colour association | Stronger than colour association | NA |  |  |
|  | f(ISL) | 6 | 5 | 5 | 5 |  |  |
|  | f(NOS) | 3 | 5 | 2 | 2 |  |  |
| Location | Value | Outside location | Mental space | NA |  |  |  |
|  | f(ISL) | 9 | 8 | 1 |  |  |  |
|  | f(NOS) | 3 | 12 | 0 |  |  |  |
| Location (specified) | Value | Letter-adjacent | Letter-overlapping | NA |  |  |  |
|  | f(ISL) | 6 | 7 | 5 |  |  |  |
|  | f(NOS) | 2 | 6 | 5 |  |  |  |
| Shape | Value | Letter | Discernible shape | Totality of colour | Aura | None | NA |
|  | f(ISL) | 7 | 3 | 2 | 2 | / | 4 |
|  | f(NOS) | 8 | 1 | 1 | 3 | 4 | 0 |
| Automaticity | Value | Willful | Automatic | Varied | Semi-automatic | NA |  |
|  | f(ISL) | 10 | 5 | / | / | 3 |  |
|  | f(NOS) | 1 | 8 | 3 | 2 | 0 |  |

A chi-square test of independence, found no significant difference between the ISL and NOS groups for the experiential category *location:* χ2(1, N = 32) = 2.418, *p* = 0.120. Cramér’s V measure of association, ϕc, was 0.275.

##### Location (specified)

Within this category, we found a large degree of heterogeneity for both groups when describing the precise location of their colour experiences. In reports where this information was available, approximately half of the ISL participants (7 out of 13) described their colour experience as overlapping with the grapheme, the others (6 out of 13) described it as occurring near or around the grapheme.

Within the NOS group we observed two cases (N = 7) of *letter-adjacent* colour experience, and five cases of *letter-overlapping* colour experiences (i.e., the colour experience was described as occurring precisely over the grapheme). However, for the remaining NOS participants we identified an additional experiential category that was not present in the ISL group, in which the location of concurrent experience could not be discerned, because the colour experiences were tied to vague descriptions that occurred in the participant’s mental space. These participants described their synaesthetic experience as containing a component (N = 6) or consisted solely (N = 1) of propositional knowledge of the grapheme-colour association. In other words, the participants *knew* the concurrent colours without being able to localise them in space [10; 43]. Finally, one participant (NOS-04), reported experiencing an embodied connection to the concurrent colour, such that she experiences her concurrent through her body. A chi-square test of independence, found no significant difference between the ISL and NOS groups for the experiential category *location (specified):* χ2(1, N = 21) = 0.257, *p* = 0.612, Cramér’s V measure of association, ϕc, was 0.111.

On first inspection reports from both groups appear to support the associator/projector subtype distinction. However, a more detailed examination of reports from the NOS group reveal a more complicated picture. Four aspects of reported experience challenge this distinction. First, unlike classic descriptions [42], in the NOS group we observed a number of individuals in which both associator and projector phenomenology co-occurred (8 out of 15) (for a similar finding, see [36]). For example, consider the following report:

> Participant NOS-12: … The colours become even more intense if I’m tired. If I’m super caffeinated. And I can become more of a projector and less of an associator during those experiences. Like if my brain is frazzled, all of a sudden, I am a projector, and not an associator.

This duality is quite apparent in Participant NOS-08. On the one hand, her synaesthetic experiences may take the form of an awareness of colour inside her mental space:

> Participant NOS-08: They are in my head. They are sort of in the middle. The colours for the alphabet, they, or with words, they are sort of in the middle of my head there but down. So, behind my eyes, and in the middle of my brain. And how I see it, it’s all black in my brain. It’s like black, with just the words in the middle inside of my head. And I’m reading it, sort of, left to right. I see them as I would on the page.

However, under specific circumstances (in this case reading), these colour experiences are reported as occurring in external space:

> Participant NOS-08:1 am basically sitting, this is my son’s bedroom, these are his bookshelves. Some of his bookshelves. And I am actually looking at some words here, all the time. And so, [pause] I am looking at *Civil War* – He’s a historian – [laughs] where it says *Civil War*, obviously I can see it written in red. But then in my head, it’s going green, white, brown, white, white, it’s going green, yellow, brown. In my head. It’s almost like I can read it. Obviously, I can see it. But it’s not changing in front of me. It’s not changing on the book spine. It’s in my head. I translated it into colour.
>
> Researcher: Yeah. […]. Are you aware of these two separate colours of the actual red of the title and then of your synaesthetic experience, are you aware of them simultaneously or do you switch between the two?
>
> Participant NOS-08:1 switch. I sort of, I can, oh, no, actually, when I am looking at it there, it’s going green, the C is going green in my head as I am looking at it there. Obviously, I’m aware of the red, and I’m reading it in red, but then it’s going into colour in my head. I know that sounds so bizarre.

Conversely, participant NOS-03 reports that when external stimuli appear spontaneously or take him by surprise, his synaesthetic experience was described as occurring in the *outside world*, but when elicited in the artificially constrained environment of the interview, his concurrent experience was described as residing within his mental space.

Second, for some NOS participants (7 out of 15) the synaesthetic experience contained a component (6) or consisted solely (1) of propositional knowledge of the grapheme-colour association. In other words, these participants *knew* the associated concurrent colour without being able to localise it in space. Others have described this instantiation of GCS as ‘know-associator’, that is, such synaesthetes simply know the colour associated with each letter but do not have a visual representation of it [36,43]. This non-perceptual subtype, does not fit within the subtype classification (even though these participants display consistency for letter-colour pairings), and may reflect a weak form of synaesthesia, in which, the perceptual concurrent experience has faded over time, but the associative aspects remain.

Third, within cases that described their grapheme-colour associations as propositional knowledge, five out of the six NOS participants described their concurrent phenomenology as consisting of multiple streams of experience. Rather than being aware of a concrete colour, the participants experienced a vague awareness or a feeling of a colour. This finding is in line with previous research, which also found that some synaesthetes could not easily describe their concurrent experience [36]. Again, these descriptions present difficulties when attempting to place them within the projector-associator continuum.

Finally, one participant (NOS-04), reported experiencing an embodied connection to the concurrent colour. Her description implies an active participation in attending to the colour such that she experiences her concurrent through her body (matching what has been referred to as *attentional dispositions*, [34,44] and *existential orientation* [45] in phenomenological literature; and *overarching states of mind* in neuroscientific literature [46]. Again, this description falls outside of the classical subtype classification.

Together, the results of the experiential categories *location* and *location* (*specified*) demonstrate that similar to NOS, ISL experiences were reported to occur either in external space (*projector*), or within their “mind’s eye” (*associator*). In terms of the specific location of colour experience (*location* (*specified*)), the majority of both groups described them as occurring either adjacent to, or overlapping with the letter. However, within the NOS group, we discovered an additional category of experience, in which the location of concurrent experience could not be discerned. A detailed examination of reports from the NOS group for these categories revealed a number of aspects of experience which are at odds with the subtype classification. Results from these experiential categories, combined with previous work [11,36,47], suggests that while initially useful as a tool for dividing synaesthetes into subgroups, the project-associator distinction may require evolving into a more nuanced phenomenological space.

#### 3.2.3 Shape of colour experience

This experiential category describes the shape or form associated with the colour experience reported by participants. All of the ISL and the majority of the NOS group described the same three forms of colour experience for this category, reporting that either their colour experience mirrored the shape of the grapheme (ISL: 7 out of 14; NOS: 8 out of 15) (coded as *letter*), that their associated colour experience had a *discernible geometric shape* (e.g., a block of colour) (ISL: 3 out of 14; NOS: 1 out of 15), or that their colour experience exhibited a shapeless presence of colour or an *aura* (ISL: 4 out of 14; NOS: 6 out of 15)(the participants’ entire mental space was filled with the awareness of the associated colour)(see Table 4). In terms of the specific shape of synaesthetic colour experience, we identified two additional experiential categories within the NOS group. Specifically, a distinction was established between cases where colour experiences where explicitly described as having no shape (coded as *none*; N = 4) and a single case in which the participant described the colour experience as taking up the whole of their mental space (*totality of colour*), with it becoming part of their sense of embodiment. We note that values for the category *shape* may overlap; i.e., an individual may associate one letter with a specific shape and another letter with a different shape. Thus, we observed one case where the shape of the letter co-occurred with the shape of an aura, and one case where it co-occurred with totality of colour. A chi-square test of independence, found no significant difference between the ISL and NOS groups for this experiential category: χ2(2, N = 22) = 1.228, *p* = 0.541. Cramér’s V measure of association, ϕc, was 0.236.

#### 3.2.3 Relative strength of colour experience

This category refers to a specific mental exercise participants were asked to perform during the interview, in which they were asked to compare the vividness of their strongest trained/natural synaesthetic colour experience to visual imagery associated with a life-long colour association of a real-world object (e.g. the specific shade of red associated with an English post-box). We created three values to describe the strength of synaesthesia-like experience: *weaker than non-synaesthetic colour association, stronger than non-synaesthetic colour association*, and *equal to non-synaesthetic colour association*. However, we note that the strength values of synaesthesia-like experience may overlap, as we found that experiences in both groups displayed a high degree of heterogeneity from grapheme to grapheme.

Following the mental exercise, the majority of participants in both groups reported their associated synaesthetic/synaesthesia-like colour experience as being equal to or weaker than visual imagery associated with a non-synaesthetic colour association (ISL: 11 out of 16; NOS: 11 out of 15). Five ISL and two NOS participants reported that their synaesthetic experience was stronger than visual imagery associated with a life-long colour association (see Table 4). Interestingly, NOS participants frequently reported that there was no qualitative difference between their synaesthetic experience and visual imagery of a real-world colour association, for example:

> Researcher: *Bananas are yellow* is okay. Compare that colour association with *red is rusty brown*, or sorry, *R is rusty brown*. […] How are these two statements similar or different in your experience?
>
> Participant NOS-10: They are pretty much exactly the same, to be honest. [pause] There’s no difference. Bananas will always be yellow, and R is always going to be rusty brown.

Findings for this experiential category may be interpreted as suggesting an equivalence between visual imagery and synaesthetic experience. Indeed, ever since synaesthesia was first described, debate has continued over whether synaesthetic experiences should be viewed as distinct from vivid mental imagery [48,49,50]. Some synaesthetic experiences are described as occurring within the “mind’s eye”, which is evidently similar to descriptions of visual mental imagery. Nonetheless, phenomenological differences exist between these two forms of perception. For example, in projector variants of GCS, the concurrent experience is described as being externally localised, whereas visual imagery is normally reported as occurring within the “mind’s eye”. In addition, visual imagery is also associated with a sense of volition, whereas a widely agreed hallmark of the synaesthetic concurrent experience is that they are automatically triggered by specific inducing stimuli [37]. We will return to this point shortly.

#### 3.2.4 Automaticity of colour experience

This category of experience refers to whether the colour experience was described as being *automatic* (i.e., experienced as if it happened to the participants) or *wilful* (i.e., experienced as if it was performed by the participants). In our previous training study, we reported that 12 out of 18 ISL participants indicated that the translation from letter to colour did not require any effort, 3 out of 18 were undecided and 3 out of 18 said the translation required effort [2]. In contrast, to these findings, our qualitative analysis revealed that for the majority of ISL participants (9 out of 14) their synaesthesia-like experience required mental effort to occur, with only five cases reporting that the experience was automatic. A possible explanation for this divergence relates to the different prompts (open-ended vs. closed-form) used to assess automaticity, which may have led participants to provide an answer based on different aspects of their experience [e.g.51,52]. In our original training study automaticity was assessed by asking participants: ‘Does it require mental effort from your side, in order to have these experiences?’. By contrast, the results of this experiential category were based on an analysis of each participants entire interview (data here https://osf.io/e367d/?view_only=dd61d42daa7a4c848023b89bd38789f8).

Using the same qualitative analysis, in contrast to the ISL group, we found that the majority of NOS participants (8 out of 15) described their synaesthetic colour experience as fully automatic (see Table 4). This result was confirmed using a forced choice question, in which participants rated the automaticity of their synaesthetic experience from 1-10, 10 being fully automatic; the average score across all participants was 8.99 (N = 15, SD = 1.62). We found that only one participant described their synaesthetic colour experiences as being *wilful*, i.e. they needed to be wilfully brought to the front of their awareness.

This divergence in experience between the two groups was highlighted by a chi-square test of independence, which identified a significant difference between the ISL and NOS groups for the experiential category *automaticity:* χ2(1, N = 24) = 4.934, p = .026. Cramér’s V measure of association, ϕc, was 0.453. In reference to the category *stability of experience*, it needs to be restated that the automaticity of letter-colour associations varied from grapheme to grapheme.

While the ISL group only reported on the distinction between *wilful* and *automatic* experiences, we identified three additional aspects of NOS experience for this category. The first is *contextually varied experience*, which describes individuals whose synaesthetic experiences displayed different levels of automaticity based on their immediate situation. For example, we found three NOS participants who reported that while reading prose, were able to ignore their synaesthetic experiences, and three participants whose synaesthetic colours were dependent on the context in which they were viewed.

The second additional experiential category identified within the NOS group is *semi-automatic experience*, which refers to situations in which participants experience the option of bringing their concurrent experience to the forefront of their awareness or to block it from occurring at all. We observed two cases of individuals who reported *semi-automatic* experiences. Consider the following report:

> Participant NOS-04:I don’t focus in on the colour quite as much as if I am focusing on it. So, to say, like, for example, I am reading a novel. I don’t focus on the colours that are there. So, they become sort of pushed back out of my conscious thought until I individually think of a letter…. It’s a background noise type thing.

The third additional experiential category we identified is named *reflective association*. This category describes reports of colour experiences only become present in the participants’ awareness when they consider a given grapheme, or when they reflect on the letter itself. If they do not assume this reflective stance, no synaesthetic experience occurs. Surprisingly, similar to reports from the ISL group that their colour experience required mental effort, we found codes referring to the *reflective* nature of concurrent experiences in the majority of NOS participants (10 out of 15).

The category *reflective association* contains two values: *phase-in* and *phase-out*. The former describes situations in which participants perform a mental gesture in order to bring the synaesthetic experience to the forefront of their awareness, as in the following example:

> Participant NOS-05: So, this, and perhaps I don’t have the right kind of synaesthesia for you. I don’t *see* the colour when I look at the word on the page. But when I envision a letter in my mind, it has a colour. Words in my mind have a colour, and certainly days of the week and numbers very strongly have a colour. But if I look at your list, I don’t see the colours when I see the letters.

As well as the following:

> Participant NOS-06: I’m only aware of it… if I’m reading, there are no colours at all, I’m just reading. If I stop for a moment and think about synaesthesia … if someone says … or if my mind drifts from the reading to synaesthesia, then I can see letters. They are kind of there all the time. I can see them if I want to. The colours are there. I can see that it’s black. It’s just black. The text. But if I want to see the colours, they will pop up in a way, but it’s different in a way, because I am physically seeing the black letters, and I am mentally seeing the colours. Because it happens so automatically – that’s it! It’s the synthesized bit of synaesthesia, in that it is so automatic that I am imagining the colour, it’s not in the imaginary visualization way, because it’s just there and I can tap into my brain or not. And most of the time, I never do.

Conversely, the value *phase-out* refers to the mental gesture whereby participants move their synaesthetic experience to the back of their mind (see excerpt from participant NOS-04, above). If they did not assume this reflective stance, no synaesthetic experience was reported.

Similar reports of mental action altering perceptual experience have also been described in other forms of synaesthesia, such as sequence-space synaesthesia. In a single case study, the concurrent experience was described as occurring within a “mental room”, with the participant reporting the ability to selectively shift their attention to either the mental room in which the concurrent occurred, or to the real world (Gould, et al., 2014). Within the NOS group we identified seven participants who reported a similar ability: while reading prose, they were able to ignore their synaesthetic experiences, pushing them to the back of their mind.

The identification of this experiential category *reflective association* suggests that, for most NOS participants, an inducer must be attended to in order to elicit a concurrent experience. This finding is in line with previous work, which suggests that attention must be deployed to a grapheme for the associated colour experience to emerge [36,53–56]. Indeed, a key feature of the description of automaticity in relation to concurrent experience is that an inducer must be sufficiently processed to elicit a concurrent experience. For example, within a visual search task, in which a target digit was presented within an array of distractors, both synaesthetes and matched-controls were shown to be equally inefficient at locating the target when the target and distractors were achromatic, despite the same digit eliciting a distinct colour experience for the synaesthetes when presented outside of the search task [54,56]. The authors suggest that synaesthetic colour experience does not arise early enough in visual processing to attract focal attention. In addition, other studies have shown that reducing the available attentional resources when processing an inducing stimulus significantly reduces synesthetic interference effects, within a modified Stroop task [53,54]. These findings collectively suggest that the concurrent experience requires attention to be focused on an inducer in order to occur. However, the precise meaning of ‘sufficiently processed’ is unclear and open to interpretation, if it refers to a participant being consciously aware of the stimulus, then our findings differ from the literature, as some participants reported being aware of a letter, but needed to further reflect on its associative aspects to elicit a concurrent experience. Based on the subjective reports of the majority of our NOS participants, the category *reflective association*, supports the notion that in addition to inducers requiring selective attention, they also need to be considered in order for them to elicit a synaesthetic colour experience. Further research is required to validate if this particular aspect of synaesthetic experience extends to a wider GCS population.

The results of the chi-square test suggests that a distinguishing aspect of experience between ISL and NOS groups is that NOS experiences are subjectively perceived to occur automatically, whereas ISL experiences are generally reported as being wilful. However, the category *reflective association* leads to an apparent contradiction in our data, namely, for NOS participants the closed-form questions point to concurrent experiences being automatic, while the qualitative data suggests that for the majority of participants they are experienced as being wilful. How can we explain this contradiction in our data? One possibility is that, similar to the ISL group, the different prompts (open-ended vs. closed-form) lead the participants to provide an answer based on different aspects of their experience [e.g. 51]. The open-ended questions (within the context of the interview) may have been interpreted as enquiring about the awareness of colour in their visual consciousness, whereas the closed-form measures may have been interpreted as referring to their propositional knowledge about the grapheme-colour association. For example, the associative link between ‘*R*’ and ‘*red*’ may be automatic in a participant’s experience, whereas the visual experience of redness elicited by the letter ‘*R*’ seems to be associated with the reflection on the letter itself.

Observations from this experiential category also speaks to the relationship between synaesthetic experience and visual imagery. As previously mentioned, a key distinction between these two forms of perception is volitional control, - a defining characteristic of synaesthetic experiences is its automaticity, while visual imagery usually exhibits an element of volitional control. However, we found that some NOS participants reported volitional control over their synaesthetic experience. Additionally, the majority of NOS participants reported that their concurrent experience only occurred following a volitional *reflective association*. We acknowledge that visual imagery is a complex and flexible phenomenon; that likely consists of many components [57]. However, reports of NOS participants in our study provides tentative evidence that for some synaesthetes their concurrent experience exhibits a degree of volitional control, which appears similar to visual imagery [58].

Finally, individual variation in the vividness of visual mental imagery in the general population has been well documented [58,59]. We note that reports from both ISL and NOS participants displayed individual variations in the vividness of their concurrent experience. This raises the question, if variations in visual mental imagery could account for individual differences in the reported vividness of synaesthetic concurrent experiences observed in this study? Indeed, grapheme-colour synaesthetes have been shown to display more vivid visual imagery compared to matched controls [60]. A finding which has led some researchers to conclude that there must be a continuum of experience in GCS, from those who never have a visual experience of a concurrent (possibly reflected by the non-perceptual subtype) to those individuals who always enjoy a vivid conscious concurrent [49]. We did not explore variations in visual mental imagery within our NOS participants; future work is warranted to examine if variations in visual mental imagery affects the reported vividness of concurrent colour experience.

To aid in the interpretation of results, Table 4, summarises the frequency distribution of phenomenological dimensions between the ISL and NOS groups.

### 3.3 Implications for synaesthesia research

Our phenomenological analysis of the NOS group raises a series of points for discussion regarding the core definitional characteristics of natural synaesthesia, which we now consider in terms of its internal consistency, automaticity and spatial characteristics [7,10,37,61].

The consistency of synaesthetic experience has become the behavioural ‘gold standard’ for determining the genuineness of the condition, being described by some as a fundamental characteristic of synaesthesia [11]. Using the colour-consistency test for the behavioural diagnosis of synaesthesia [7] our previous training studies, including the participants reported on here, found that trained graphemes demonstrated levels of consistency indicative of synaesthetic experience [1,2]. These results suggest that, while a useful indication of natural synaesthesia, consistency cannot be viewed as unique to naturally occurring synaesthesia. Indeed, debate exists around the validity of consistency as a diagnostic criterion in synaesthesia research. For example, Simner et al., [62] found that not all synaesthetes meet the criteria of consistency even though they feel strongly that they experience the condition, and also noted that some synaesthetes reported that their concurrent experiences were never consistent. Some have argued that the criterion of consistency over time provides a circular definition; that is, it fits the profile of synaesthesia described in the literature precisely because this group has been selected based on this criterion [41,63]. It is possible that studies investigating synaesthesia have been self-selecting a biased sample of consistent synaesthetes, while claiming that consistency is a defining feature of this condition.

A wealth of contemporary studies have described the automaticity of synaesthetic experiences, i.e. a concurrent experience is automatically triggered by an inducer and is not under voluntary control, as another hallmark of the condition, which is normally assessed using an adapted version of the Stroop task [4,5]. We found, using subjective reports, that the majority of NOS participants reported a need to selectively attend to the inducing stimulus in order for a concurrent experience to be elicited. This finding is in line with previous research [8,36,53,54,55,56]. However, we found that the majority of NOS participants also needed to ‘reflectively consider’ the inducer, in addition to paying attention, to elicit a concurrent experience. We also found that three NOS participants reported that while reading prose they were able to ignore their synaesthetic experiences. How do these findings fit within the wider synaesthesia literature? A key feature of the description of automaticity in relation to concurrent experience is that an inducer must be sufficiently processed to elicit a concurrent experience. However, this description is open to interpretation, if sufficiently processed refers to a participant being consciously aware of the stimulus, then our findings differ from the literature. This difference is highlighted by some participants not experiencing a synaesthetic concurrent in circumstances where this would be undesirable (e.g., reading prose), even though they are consciously aware of the inducer. If, on the other hand, sufficiently processed refers to paying close attention to the inducing stimulus (i.e., focusing attention to the stimulus and its associations), then our findings are in line with previous research.

In standard terminology, natural synaesthetes have often been characterised along a projector to associator continuum [3]. In both ISL and NOS groups, some participants indeed reported that their concurrent occurred within their mind’s eye, while others indicated that it appeared in external space. However, we found that some NOS participants reported both types of description, while others stated that there was no specific spatiality associated with their concurrent experience. Therefore, while our results suggest that some synaesthetic experience may be spatially localised, we found other variants that lacked a defined spatial location. Our findings suggest that there is no clear rationale for future studies to subdivide synaesthetes simply based on the spatial location of their concurrent experience.

The comparisons between induced and natural synaesthesia described in this paper cast further doubt to the claim that natural synaesthetic phenomenology can only occur in a rare subset of the population that exhibit a genetic predisposition [38,64,65]. Even in cases where a genetic predisposition leads to the development of synaesthesia, the observation that in most forms of synaesthesia concurrent experiences are triggered by cultural artefacts, suggests that natural synaesthesia must include a substantial dependence on learning and prior experience [1,2,13,66,67].

### 3.4 Limitations and future directions

The phenomenological descriptions reported here are by no means exhaustive, particularly with regards to naturally occurring synaesthesia (NOS). Indeed, even within the present set of transcripts, additional experiential themes and categories were identified, which remain to be fully explored.

Among these additional categories, *veridicality*, is an aspect of experience that describes if participants colour experience lacked either veridicality or what has been called “perceptual presence” i.e. they were not perceived as properties of the world [68]. The majority of both groups provided similar descriptions about the veridicality of their associated colour experiences, which was compatible with a lack of perceptual presence (for more detail see supplementary material https://osf.io/e367d/?view_only=dd61d42daa7a4c848023b89bd38789f8). There were possible exceptions, but the evidence we have speaks more to the location of concurrent experience rather than to its continuity with ‘real world’ perception in the sense of perceptual presence. For example, in one surprisingly expressive example, participant ISL-12 describes their experience as follows: “I would say that they feel roughly the same, but it might sound stupid, but like tomato is more of a real colour, but when I think of the green (trained colour) I think of it on a computer screen and it’s a sort of not a real thing, so it doesn’t feel as natural. When you think of a tomato you think about how the light hits something, that shape, and that’s part of the colour”. This single quote nicely illustrates how this participant experiences appear to lack perceptual presence. Further research is needed to examine if induced or natural synaesthetic experience do indeed lack perceptual presence and what cognitive and neural underpinnings may explain this aspect of experience compared to normal, non-synaesthetic perception.

Another identified experiential category within the NOS group was *functionality*, which describes whether individuals can use synaesthesia as a beneficial cognitive strategy, or whether the synaesthetic experience is disruptive (see annotated codebook for further description). As well as the widely reported cognitive benefits found in natural synaesthesia, such as an enhanced memory [for a review, see 69], we found that for our participants any benefits co-occurred with detrimental effects, such as interference effects in daily life between the concurrent experience and inducer.

In addition, we identified the experiential category *colour-as-a-feature*, which describes a common report within the NOS group, whereby the colour is understood as a basic, elementary feature of the inducing stimulus, similar to its shape or the phoneme associated with a given letter. Further research is required to identify if this is a general feature of GCS.

The interview also included a detailed exploration of how the phonetics of a grapheme, its outward appearance or its underlying meaning affected the concurrent experience. For example, some participants reported that visually distinct forms of a grapheme induced the same colour, as long as they were members of the same linguistic category, i.e. ‘b was blue’ for all instances of the letter ‘b’ regardless of its font or even when presented in other non-Roman languages (e.g. the letter Б in Cyrillic). These data could be used to further explore the relative contributions of conceptual associations or the sensory/perceptual features of a grapheme in the generation of synaesthetic phenomenology [70,71].

The phenomenological dimensions identified here display a large degree of overlap with previous reports of synaesthesia (e.g. 3,11,37]. It is therefore likely that these dimensions are typical of a wider grapheme-colour synaesthesia (GCS) population. However, we acknowledge the conclusions drawn in our study are limited to our participants and the training regime used [2]. Further work is required to examine if the identified experiential overlap between induced and naturally occurring synaesthesia apply to a wider population. In addition, we acknowledge that qualitative analysis is particularly susceptible to researcher bias. We employed a number of strategies to mitigate this influence, such as the annotated codebooks and intercoder verification. We also provide access to all interview data as raw transcripts, alongside the annotated codebooks and saturation grids (see supplementary material https://osf.io/e367d/?view_only=dd61d42daa7a4c848023b89bd38789f8) However, we acknowledge that qualitative research does not reflect an objective, opinion-free point of view, but we suggest that its strength is in adding a unique level of understanding to a condition such as synaesthesia, which in turn can be used to drive future experimental work.

Demographic information from the NOS group revealed that six out of the fifteen participants worked in mind-science areas (e.g. Psychology, Neuroscience), raising the possibility that they may have provided theoretically-laden descriptions of their experience. NOS represents a relatively small subset of the population, with many researchers relying on the same group of synaesthetes across multiple studies. Future studies should take care to make sure that participants detailed knowledge of the synaesthesia are not influencing their results, creating a circular confirmation of the condition; that is, the profile of synaesthesia described in the literature occurs precisely because participants are aware of normative responses within NOS.

A potential limitation of our training studies is the possible influence of demand characteristics – i.e., the social dynamics present in a researcher-participants relationship, whereby the latter modifies their behaviour in accordance with what the participants perceive to be the research goal of the study [72]. To guard (as far as possible) against this potential influence, the term ‘synaesthesia’ was not referred to at any point during the study. The training protocol was always described as a memory training task. The interview, however, was the first point in the study where a possible connection to synaesthesia was alluded to by the researcher. Following completion of the study a debriefing email was sent to all subjects asking if they had become aware of the synaesthesia component of the study at any point (yes/no responses only). 15/18 subjects answered no, indicating that for most participants demand characteristics were minimal, if present at all. This combined with training induced behavioural and neurophysiological effects argue against demand characteristics driving responses during the interview [2]. However, we acknowledge the potential confounding effects of demand characteristics in driving experience within psychological science, especially for subjects who score high on scales of hypnotic suggestibility or ‘phenomenological control’ [73]. Phenomenological control refers to an individual’s ability to alter what they experience, both within and outside of the hypnotic context, in ways that are consistent with their plans and goals [74,75]. Recent work has investigated the possible influence of phenomenological control in generating experiences in mirror-sensory synaesthesias: mirror touch, in which the observation of someone being touched elicits a reported tactile sensation in the observer [76]; and vicarious pain, in which observed pain elicits reports of experienced pain [77]. Lush et al., [74] found that hypnotisability scores strongly predicted mirror-sensory synaesthesia responses, suggesting that some of the experienced touch and pain sensations may have been the result of the participants’ capacity to ‘create’ the experiences of touch using phenomenological control. The authors suggest that in the case of mirror-sensory synaesthesias, individuals may habitually (but involuntarily) implement phenomenological control in everyday life, when it is in-line with their goals, creating tactile sensations. These findings raise the possibility that habitual phenomenological control may also underlie the reported colour experiences in both induced and natural synaesthesia. To address this question, future studies investigating phenomenology within grapheme-colour synaesthesia should also administer measures of phenomenological control such as the Sussex-Waterloo Scale of Hypnotisability (SWASH; [78]), to investigate if synaesthetic phenomenology is related to individual differences in phenomenological control ability.

Finally, extensions of our approach may be used to advance neurophenomenological accounts of synaesthesia. For example, first-person data could be used as a heuristic to describe and quantify the neurodynamics associated with synaesthesia. Indeed, some have already shown how individual differences in synaesthetic phenomenology (localisation) are associated with characteristic neural responses [79]. It remains an open question to what extent associative training paradigms that lead to dramatic alterations in phenomenology, such as those described here, can modify structural and large-scale dynamical features of the human brain.

## 4.0 Conclusion

We report the results of an in-depth qualitative analysis of a specific form of training-induced synaesthesia, resulting from extensive and adaptive associative training, in conjunction with an analysis of responses to a similar analysis performed on naturally occurring grapheme-colour synaesthetes. We compared commensurate categories for both forms of novel perceptual experience, identifying 5 main experiential categories that were common across induced and naturally occurring synaesthetic experience:

1. Stability of colour experience.
2. Location of colour experience.
3. Shape of colour experience.
4. Relative strength of colour experience.
5. Automaticity of colour experience.

For these experiential categories only reports relating to the automaticity of colour experience differed significantly between the two groups, with NOS experiences being described as mostly automatic, whereas induced ISL experiences were mostly described as being ‘wilful’. However, descriptions of automaticity differed within the ISL and NOS groups depending on the method of questioning used (open-ended vs. closed-form). Together, these results suggest that as with other experiential categories, this aspect of synaesthetic experience displays a high degree of heterogeneity, which further varies depending on the method of questioning.

While many of these experiential categories have been identified elsewhere [3,11,37], previous work has provided only limited information about the first-person experience of the participant in relation to each phenomenological category. In contrast, the present study provides an extensive description of the phenomenology of induced and natural synaesthetic experience, which has revealed a surprising degree of heterogeneity in the experience, both within and between individuals experience for all experiential categories. This rich resource can be used as a basis for devising future investigations.

Finally, our results extend previous reports of the parallels between induced and natural synaesthetic experience [1,2]. While we refrain from ascribing a phenomenological equivalence between training-induced and natural synaesthetic experiences, in combination with our previous findings [1,2], our results provide strong evidence that intensive and adaptive training of letter-colour associations can alter conscious perceptual experiences of non-synaesthetes, producing a coordinated set of synaesthesia-like characteristics, which bear striking similarities to those found in natural synaesthesia.

## Acknowledgments

All authors are grateful to the Dr. Mortimer and Theresa Sackler Foundation, which supports the Sackler Centre for Consciousness Science. AO acknowledges the European Commission for support through the ERASMUS+ scholarship. AKS acknowledges additional support from the Canadian Institute for Advanced Research (CIFAR) via their Azrieli Programme on Brain, Mind, and Consciousness. DB is funded by Wellcome Trust grant 210920/Z/18/Z. We are grateful to Carys Barnfield, Acer Y. C. Chang, Elena Gelibter and Alex Piletska for assistance with data collection in Rothen et al. (2018). We would also like to thank Heather M. Iriye for her help with the processing of the Stroop data for Rothen et al. (2018).

